# Thermostabilization and purification of the human dopamine transporter (hDAT) in an inhibitor and allosteric ligand bound conformation

**DOI:** 10.1101/335968

**Authors:** Vikas Navratna, Dilip K. Tosh, Kenneth A. Jacobson, Eric Gouaux

**Affiliations:** Vollum Institute, Oregon Health & Science University, Portland, Oregon, United States of America; Molecular Recognition Section, Laboratory of Bioorganic Chemistry, National Institute of Diabetes and Digestive and Kidney Diseases, National Institutes of Health, Bethesda, Maryland, United States of America; Howard Hughes Medical Institute, Oregon Health & Science University, Portland, Oregon, United States of America

**Author notes:** Corresponding author (EG).

## Abstract

The human dopamine transporter(hDAT) plays a major role in dopamine homeostasis and regulation of neurotransmission by clearing dopamine from the extracellular space using secondary active transport. Dopamine is an essential monoamine chemical messenger that regulates reward seeking behavior, motor control, hormonal release, and emotional response in humans. Psychostimulants such as cocaine primarily target the central binding site of hDAT and lock the transporter in an outward-facing conformation, thereby inhibiting dopamine reuptake. The inhibition of dopamine reuptake leads to accumulation of dopamine in the synapse causing heightened signaling. In addition, hDAT is implicated in various neurological disorders and disease-associated neurodegeneration. Despite its significance, the molecular architecture of hDAT and its various conformational states are poorly understood. Instability of hDAT in detergent micelles has been a limiting factor in its successful biochemical, biophysical, and structural characterization. To overcome this hurdle, first we identified ligands that stabilize hDAT in detergent micelles. Then, we screened ∼200 single residue mutants of hDAT using high-throughput scintillation proximity assay, and identified a thermostable variant(I248Y). Here we report a robust strategy to overexpress and successfully purify a thermostable variant of hDAT in an inhibitor and allosteric ligand bound conformation.

## Introduction

Neurotransmission is initiated by the release of neurotransmitters from the pre-synaptic neuron into the synapse. The SLC6 family of secondary active transporters such as dopamine transporter(DAT), norepinephrine transporter(NET), and serotonin transporter(SERT) play a crucial role in the regulation of neurotransmission(1). These monoamine transporters, also known as neurotransmitter sodium symporters(NSS), are located on the plasma membrane of a pre-synaptic neuron and terminate the neurotransmission. The termination is achieved by returning the neurotransmitters from the synapse to the pre-synaptic neuron, thereby maintaining neurotransmitter homeostasis at the chemical synapses. This process, known as reuptake, begins with the binding of the neurotransmitter to a sodium-ion-primed outward-open conformation of the transporter. The transporter then undergoes a series of conformational changes to assume an inward-open conformation thus returning the neurotransmitter to the pre-synaptic neuron. Then, the transporter reorients itself to an outward-open conformation and the returned neurotransmitter is either sorted into a vesicle for reuse or is degraded(2-5).

Structural characterization of the bacterial orthologues of NSS members, such as LeuT and MhsT, provided initial insights into the molecular basis of these conformational transitions (6-17). However, the structures of these bacterial orthologues do not provide a basis for the diverse pharmacological profile, ligand specificity, and allosteric regulation displayed by the eukaryotic NSS members. The human dopamine transporter(hDAT) plays a crucial role in dopamine homeostasis at the dopaminergic neurons. In the past decade, over twenty coding variants of human dopamine transporter(hDAT) have been discovered. Some of these variants result in a dysfunctional hDAT, and are implicated in attention deficit hyperactivity disorder (ADHD), bipolar affective disorder, autism spectrum disorder, anxiety, addiction, autism, schizophrenia, Parkinson’s disease, and depression(18-20).

hDAT is the major target of addictive psychostimulants such as cocaine, which bind to the active site and prevent the conformational transition of the transporter, thereby inhibiting the reuptake of dopamine(DA)(4, 21, 22). In addition to these potent inhibitors, certain(N)-methanocarba nucleoside analogs(MRS compounds, named after the Molecular Recognition Section of NIH) interact with hDAT and prevent transport(23, 24). The nucleoside analogues were initially derived from A_3_ adenosine receptor(AR) agonists, and were subsequently structurally modified(Fig S1a) to enhance the interaction with DAT. Some of these AR agonist analogs have been reported to modulate the binding of tropane based cocaine analogs to hDAT, suggesting that they do not compete with inhibitors for the central binding site. The ability of these compounds to modulate the uptake and the ligand binding properties of hDAT without competing for the central binding site makes them novel allosteric agents. Molecular characterization of an interaction between allosteric ligand and hDAT will enable the development of therapeutics that target less characterized sites on the transporter.

Expression, purification, and structure determination of recombinant eukaryotic membrane proteins are challenging tasks. Several strategies have been developed to enhance the expression of recombinant membrane proteins, noteworthy of which is the novel recombinant baculovirus-mediated eukaryotic protein expression system. The baculovirus expression system contains genetic elements that allow efficient transcription, RNA splicing, and mRNA processing, making it suitable for the expression of low yielding eukaryotic proteins in mammalian cells(25, 26). Development of a wide variety of detergents, along with strategies such as antibody fragments, engineering of target protein-T4 lysozyme fusions, and usage of lipidic cubic phase for crystallization have aided in successful purification and crystallization of several membrane proteins(27-30). However, the most promising strategy for G-protein-coupled receptors and NSS members has been the use of conformation-specific thermostabilizing site-directed mutations. Mutations that enhance thermostability often also increase yields during purification, resulting in enhanced production of homogenous and stable protein for crystallization(31-35).

Here we describe an efficient strategy for expressing and purifying hDAT. First, using the baculovirus-mediated expression system we expressed hDAT in HEK293S cells. Next, we employed MRS compounds to stabilize the transporter, and identified conditions that allowed us to extract active hDAT from cells using detergent. Finally, we generated ∼200 variants of hDAT by site-directed mutagenesis, and by employing a scintillation proximity assay(SPA)-based thermostability screening we identified a thermostable variant of hDAT. Incorporating the insights from the screening experiments we developed a robust protocol for the purification of hDAT in an inhibitor and allosteric ligand bound conformation.

## Materials and Methods

### Cloning and site-directed mutagenesis

The codon optimized gene encoding full length human dopamine transporter was synthesized by Bio Basic, Inc., was cloned into the pEG BacMam expression vector and was expressed as a fusion protein containing an *N*-terminal His_8_-GFP-tag(hDAT-FL). Single residue mutations were made in hDAT-FL background using site-directed mutagenesis. Other constructs that are described in the manuscript are:(i) thermostable mutation made in hDAT-FL(hDAT-I248Y) and(ii) *N*-terminal 56 residue deletion in hDAT-I248Y(NΔ56-I248Y). For large scale protein expression and purification, GFP in these constructs was replaced with Strep-tag II enabling affinity purification using Strep-Tactin resin.

### Cell culturing, transient transfection and transduction

The thermostability screening was performed by expressing mutants in adherent HEK293S cells as follows. Briefly, the cells were allowed to adhere onto a poly-D-lysine coated, tissue culture treated 96-well plate in Dulbecco’s Modified Eagle Medium(DMEM) with 10% fetal bovine serum(FBS) at 37 °C. Immediately before transfection, the media was replaced with 200 μL/ well of fresh DMEM(pre-warmed to 37 °C). The cells were transfected with 100 ng/ well DNA using Lipofectamine 2000(ThermoFisher Scientific) according to the manufacturer’s protocol. The transfected cells were incubated at 37 °C with 5% CO_2_ for 10-12 h, followed by an incubation of 36 h at 30 °C. Before moving the cells to 30 °C, to enhance the expression of mutants, each well was supplemented with 10 mM sodium butyrate. The media was aspirated after 36 h at 30 °C, and the cells were resuspended in the assay buffer for thermostability screening. To screen a library of potential thermostable mutants, HEK293S cells were transfected and grown in a 10 cm petri dish and membranes were harvested from the cells for thermostability assay.

For large-scale purification, hDAT was expressed by transducing HEK293S cells suspended in FreeStyle 293 expression media. Briefly, the cells(at a density of 3-3.5 × 10^6^ cells/ mL) were infected using P2 baculovirus at a multiplicity of infection of 1.5. The infected cells were incubated at 37 °C with 8% CO_2_ on an orbital shaker. After 10-12 h the culture was supplemented with 10 mM sodium butyrate and was then transferred to a shaker at 30 °C. The cells were harvested 36 h after infection and were resuspended in a buffer(100 mL/cells from 6 L culture) containing 40 mM HEPES, pH 8.0, 200 mM NaCl, and 10% glycerol. The resuspended cells were either frozen in liquid nitrogen and stored at -80 °C until further use or were processed further for membrane isolation.

### Filter binding assay

The binding of radioligands to membranes expressing hDAT was confirmed by performing filter binding assays. Binding reactions were initiated by mixing HEK293S membranes expressing hDAT with 30 nM [^3^H]WIN35428, and incubating the mixture at room temperature on an end-to-end rotator for 30 min in binding assay buffer consisting of 20 mM sodium phosphate, pH 7.8, and 100 mM NaCl. The binding was terminated by filtering the reaction mixture through 0.4% polyethylenimine(PEI) soaked glass microfiber filters(Whatman GF/B 25 mm circles) using a Millipore vacuum filtration manifold. The microfiber filters were then washed twice with 3 mL of cold assay buffer, and were dropped into 5 mL of Ultima Gold liquid scintillation cocktail(PerkinElmer) for counting. Non-specific binding was estimated by including 100 μM GBR12909 in the reaction mixture. All binding experiments were performed in triplicate and the data was analyzed using GraphPad Prism. The data from the [^H^3]WIN35428 saturation binding assay with or without MRS7292 were fit according to a single site binding model.

To compare the effect of MRS7292 on ligands that bind at the central binding site, 100 μL of reaction mixture containing 30 nM hDAT membranes and 30 nM [^3^H]WIN35428 was incubated at room temperature for 30 min. The incubation was stopped by diluting the reaction mixture to 1 mL using dilution buffer(same as binding assay buffer with or without 5 μM MRS7292), and was filtered through glass microfiber filter. Sets of identical reactions were filtered at varying time periods(0-60 min) post dilution. The decrease in counts from [^H^3]WIN35428 bound to the central binding site upon dilution with or without MRS7292 was compared by plotting total counts observed versus time after dilution.

### Scintillation proximity assay(SPA) based thermostability screening

SPA-based mutant thermostability screening was performed using transfected HEK293S cells in 96-well plates(36, 37). Cells(∼60,000) from each well were resuspended in 60 μL of buffer containing 20 mM sodium phosphate, pH 7.8, and 100 mM NaCl, and 17 μM MRS7292. To this suspension 10 μL of 300 nM [^3^H]WIN35428 was added and the plates were incubated for 30 min at room temperature. Following the incubation, 5 μL of 20 mg/ mL Cu-Ysi SPA bead suspension and 25 μL detergent stock solution(10 mM lauryl maltose neopentyl glycol(LMNG), 1 mM cholesteryl hemisuccinate(CHS), 0.2% bovine serum albumin, 20 mM sodium phosphate, pH 7.8, and 100 mM NaCl) were added to the samples. The reaction mixture also included protease inhibitor cocktail(a final concentration of 1 mM phenylmethane sulfonyl fluoride(PMSF), 0.8 μM aprotinin, 2 μg/ mL leupeptin, and 2 μM pepstatin A). [^3^H]WIN35428 binding to various hDAT mutants was measured using a MicroBeta Trilux scintillation detector. Non-specific binding was determined using 100 μM GBR12909. The plates were read continuously until the total counts reached a plateau(∼ 1 h), and then were heated at 30 °C for 10 min in an Eppendorf ThermoMixer C, and were read again. The subsequent rounds of heating/measurement, with a 5 °C increase per cycle, were repeated till the total counts were similar to non-specific counts.

Specific counts were determined by subtracting the nonspecific counts from the total counts. The percentage decrease in specific counts versus temperature was plotted, and the melting temperature(*T*_*m*_) of the protein was calculated by non-linear fitting of the data to a Boltzmann sigmoidal equation. The mutant thermostability screening experiments were performed in triplicate. The total counts observed at room temperature were considered to be an indicator of expression, and were plotted against *T*_*m*_ of mutants to generate a scatter plot. The mutants situated farthest from the origin on the diagonal were considered to be both better-expressing and thermostable. To compare the *T*_*m*_ of hDAT in the presence of various MRS compounds, and also to reaffirm the *T*_*m*_ difference between hDAT-FL and thermostable mutants, a thermostability assay was performed using membranes isolated from large scale culture of either adherent or suspension HEK293S cells. Following membrane isolation, the subsequent steps of thermostability assay were same as the procedure reported for SPA-based mutant screening.

### Saturation binding assay

To determine the effect of MRS7292 on the affinity of hDAT to [^3^H]WIN35428, we performed ligand binding assays on isolated membranes. Membranes expressing hDAT were incubated with varying concentrations(0-120 nM) of [^3^H]WIN35428 in 1mL of binding assay buffer(20 mM sodium phosphate, pH 7.8, and 100 mM NaCl) with or without 10 μM MRS7292. The reactions were incubated for 30 min at room temperature and were terminated by filtering through 0.4% PEI soaked glass microfiber filters. The filters were washed twice with cold binding assay buffer(containing 10 μM MRS7292) and were counted in liquid scintillation cocktail.

For calculating apparent binding affinity of hDAT to [^3^H]WIN35428 and [^3^H]GBR12909 in detergent micelles, we purified hDAT-I248Y in the presence of 10 μM MRS7292 and 1 μM mazindol. Purified hDAT-I248Y(∼10 nM) was incubated with varying concentrations(0-150 nM) of radioligands and counts were measured until they reached a plateau. The counts in saturation binding assay were plotted versus concentrations of radioligand and the data was fit to a single site binding curve using the GraphPad Prism program.

### Purification

The cells expressing hDAT were sonicated and the cell debris was removed by a low speed centrifugation step(2400 x g for 10 min). Membranes were harvested from the cell lysate by ultracentrifugation(185,000 x g for 1 h), and were resuspended in a buffer containing 40 mM HEPES, pH 8.0, 400 mM NaCl, and 20% glycerol. MRS7292, GBR12909 or WIN35428, and a cocktail of protease inhibitors were added to this suspension. Equal volume of detergent solution(10 mM LMNG, 1 mM CHS, 40 mM HEPES, pH 8.0, 400 mM NaCl, and 20% glycerol) was added to the suspension. The final concentration of ligands in the solubilization mixture were 10 μM MRS7232, and 4 μM GBR12909 in the presence of protease inhibitor cocktail(1 mM PMSF, 0.8 μM aprotinin, 2 μg/ mL leupeptin, and 2 μM pepstatin A). Solubilization was carried out at 4 °C while stirring for 90 min. The detergent solubilized membrane suspension was centrifuged at 185,000 x g for 1 h and the supernatant was allowed to flow through a column packed with Strep-Tactin affinity resin, at a flow rate of 0.5 mL/ min. The packed resin was then washed with 5 column volumes of wash buffer containing 40 mM HEPES, pH 8.0, 400 mM NaCl, 0.1 mM LMNG, 10 μM CHS, 20% glycerol, 2 μM MRS7232, 2 μM GBR12909, and 25 μM 1-palmitoyl-2-oleoyl-sn-glycero-3-phosphocholine(POPC). The protein was then eluted in a buffer containing 5 mM D-desthiobiotin, 40 mM HEPES, pH 8.0, 400 mM NaCl, 50 μM LMNG, 5μM CHS, 20% glycerol, 4 μM MRS7232, 2 μM GBR12909, and 25 μM POPC.

The purified protein was concentrated to 1 mg/ mL and was loaded onto a Superdex S200 column for size-exclusion chromatography(SEC). The SEC buffer was similar to elution buffer, but without desthiobiotin and with 5% glycerol. The homogeneity and ligand binding ability of the purified hDAT were confirmed by performing fluorescence-detection size exclusion chromatography(FSEC), SDS-PAGE, and SPA experiments. For FSEC experiment, 100 μL of 30 nM protein was loaded onto a 10/300 Superose 6 column pre-equilibrated with buffer containing 0.15 mM LMNG, 20 mM HEPES, pH 8.0, and 200 mM NaCl.

## Results

### Stabilization of hDAT by allosteric ligands

Solubilization of membranes using detergent is an effective approach to extract and purify transmembrane proteins. However, this approach often destabilizes the proteins, thus requiring optimization of extraction conditions. To monitor efficient extraction, we performed FSEC using lysate obtained from detergent solubilization of HEK293 cells expressing hDAT. hDAT-FL could be successfully extracted from plasma membranes using detergent(Fig 1a). However, detergent extracted transporter displayed a loss of the ligand binding activity, suggesting instability(Fig 1b). We employed the ligand binding activity of hDAT as an indicator of stability, and proceeded to identify the extraction conditions in which the transporter retained the binding activity.

**Fig 1:**
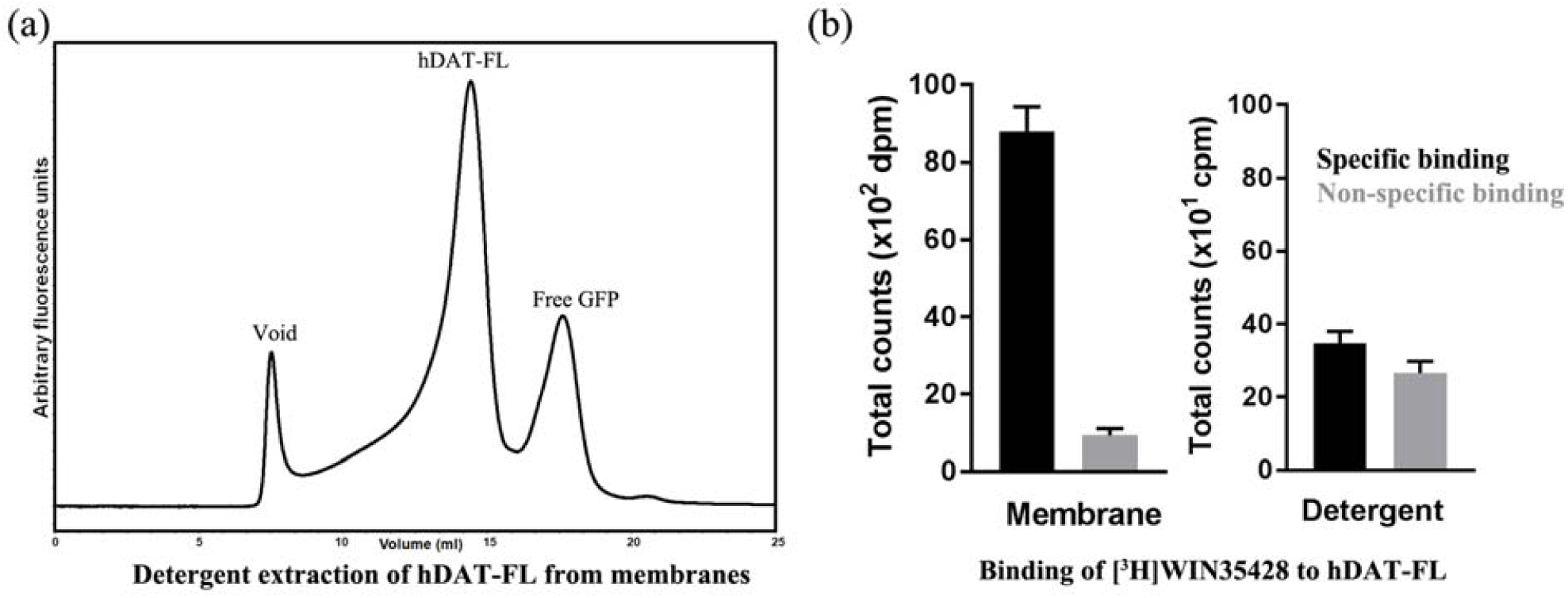
Extraction of hDAT from HEK293S cells. (a) Fluorescence of the detergent solubilized cell lysate from HEK293S cells expressing hDAT-FL is monitored by FSEC.(b) Loss of [^3^H]WIN35428 binding upon detergent solubilization of membranes expressing hDAT-WT. Shown on the left is ligand binding to membranes isolated from HEK293 cells expressing hDAT-FL, and on the right is binding to detergent extracted hDAT-FL. A representative of experiments(n=4) is shown in panel b, and the error bars denote standard errors of mean calculated from triplicate measurements.

We discovered that the ligand binding activity of detergent extracted hDAT could be preserved if the extraction was performed in the presence of(N)-methanocarba nucleoside analogs(MRS compounds)(Fig S1b)(23, 24). The MRS compounds, which contain a rigidifying bicyclo[3.1.0]hexane ring system in place of ribose, were identified as allosteric ligands of hDAT during the off-target activity screening experiments. It was reported that these conformationally-locked nucleosides interact with hDAT and modulate the dopamine transport and ligand binding(23, 24). Based on our detergent extraction experiments, it is now evident that MRS compounds can also enhance the stability of hDAT in detergent micelles. Because the *T*_*m*_ of a protein is an efficient indicator of its stability, we hypothesized that estimating the thermostability of hDAT in presence of various MRS compounds will enable us to identify most suitable MRS compound to use during detergent solubilization. To test our hypothesis, we measured the *T*_*m*_ of hDAT in the presence of different MRS compounds, and identified that MRS7292 imparted the highest degree of thermostability(Fig. 2a and Fig S1c). Capitalizing on the ability to detect ligand binding in detergent micelles in the presence of MRS7292, we designed a high-throughput SPA-based thermostability assay to discover amino acid substitutions that stabilize hDAT bound with an inhibitor, and an allosteric ligand.

**Fig 2:**
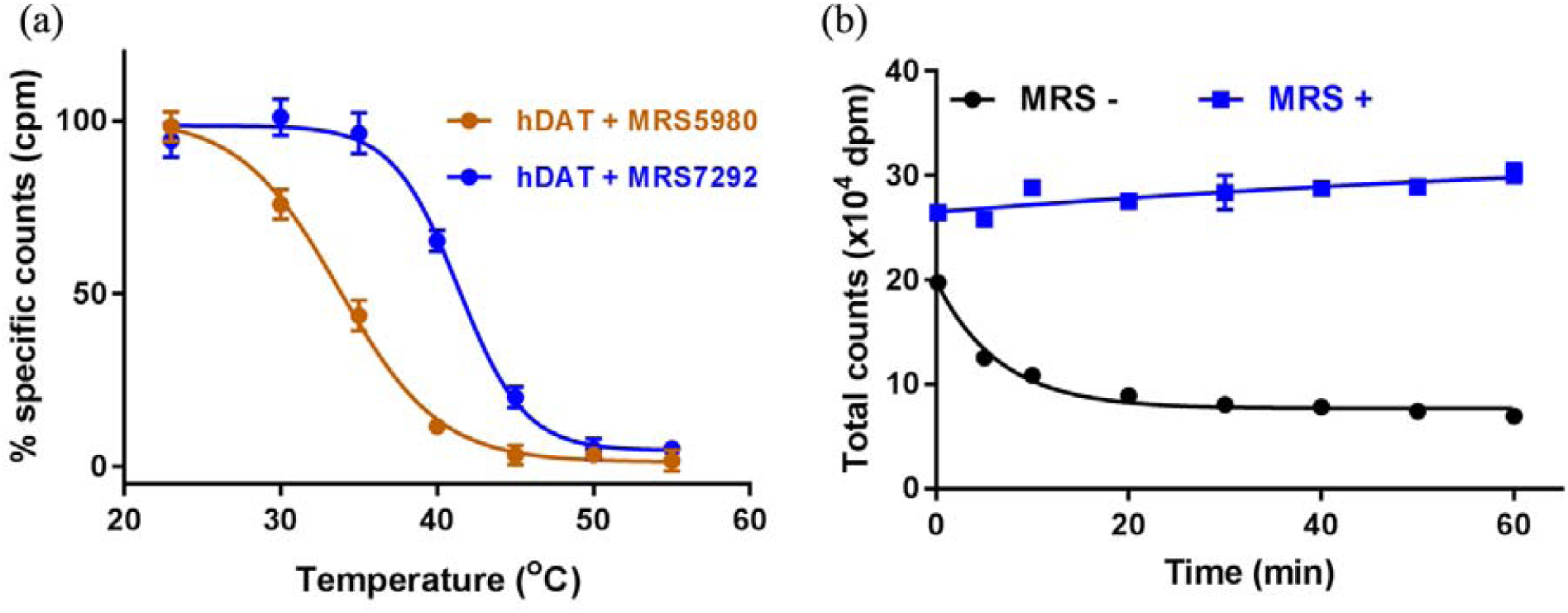
Effects of MRS7292 on radioligand binding and stability. (a) Enhanced thermostability of hDAT-WT in detergent micelles in presence of MRS compounds. hDAT bound to MRS7292(blue) displays ∼ 7.5 °C greater *T*_*m*_ than hDAT-MRS5980(red) complex, and(b) MRS7292 prevents dissociation of [^3^H]WIN35428 from the central binding site upon dilution(blue). A representative of experiments(n=3 for panel a; n=2 for panel b) are shown, and the error bars denote standard errors of mean calculated from triplicate measurements.

### Identification of sites for mutagenesis

Two approaches were employed to identify the residues to be mutated. First, a consensus sequence based site-directed mutagenesis approach was employed, in which a multiple sequence alignment was performed using protein sequences of eukaryotic DATs and prokaryotic NSS homologs from thermophilic organisms. The consensus sequence was then compared to the sequence of hDAT. The residues in hDAT that differed from the consensus were shortlisted and substituted with the consensus residue or an Ala or a Phe. In the second approach, the residues that enhanced the thermostability in dDAT and hSERT were mapped on to the sequence of hDAT, and similar substitutions were made in hDAT. The residues identified by both approaches were distributed throughout the protein. We note that residues involved in post-translational modification and zinc binding sites were excluded(Fig. 3). The His_8_-tag on hDAT allowed the monitoring of radioligand binding in SPA experiments using the Cu-Ysi SPA beads, and the GFP-tag allowed monitoring of the transfection and expression using epifluorescence and FSEC.

**Fig 3:**
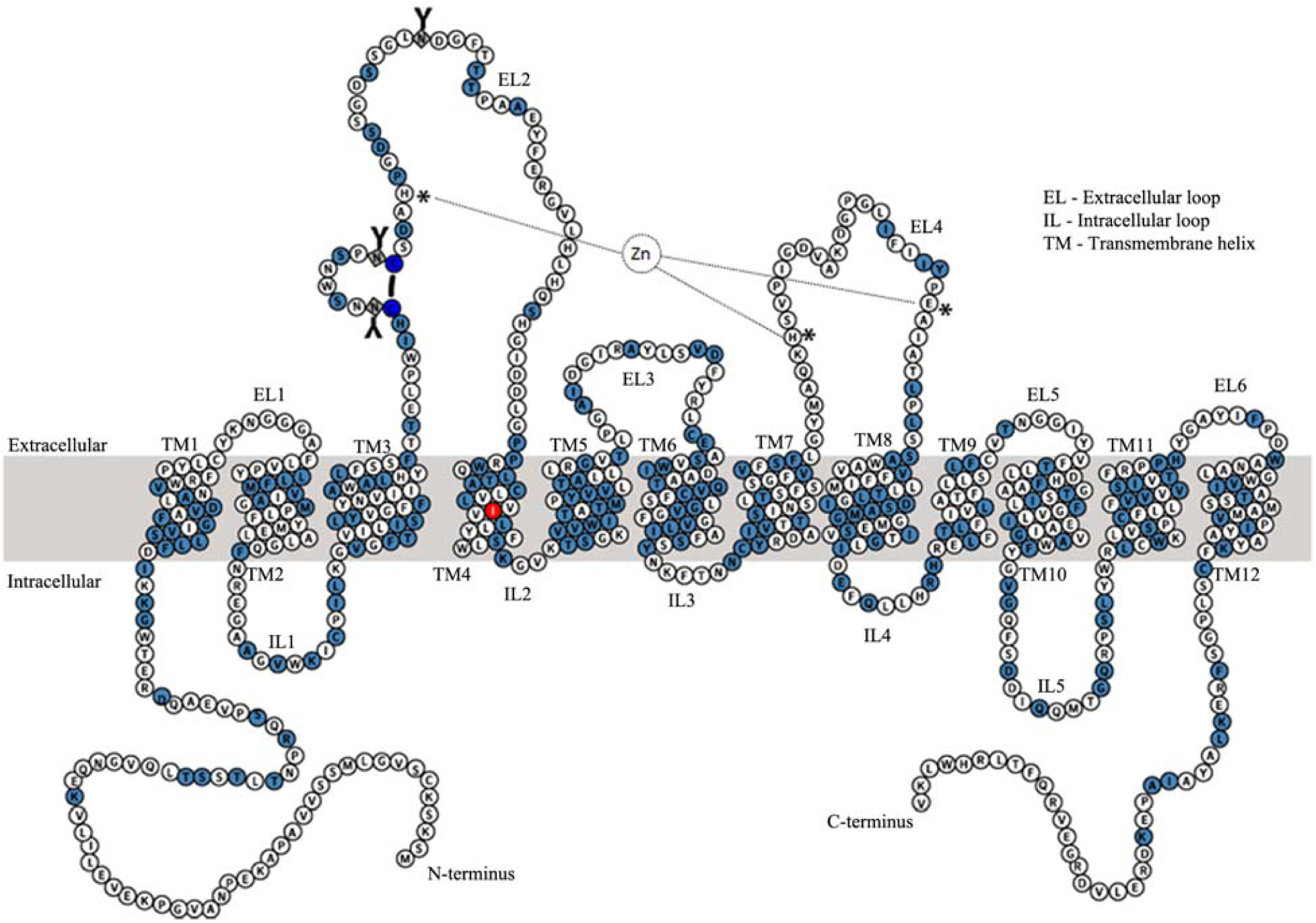
Predicted membrane topology of hDAT highlighting the mutated residues. The mutated residues are shaded in light blue. The thermostable I248Y is shaded in red. Residues comprising Zn^+2^ binding site(H193 and D206 of EL2; H375 and E396 of EL4) are highlighted by an asterisk. The black line connecting two cysteines(dark blue circles), denotes the disulfide bond between C180 and C189 in EL2. The residues(181, 188, and 205) that undergo *N*-linked glycosylation(**Y**) are represented as squares. The boundaries of transmembrane helices were defined based on an hDAT model constructed using hSERT crystal structure(5I6X) as template.

### High-throughput thermostability screening of mutants

We tailored a previously developed high-throughput SPA-based thermostability screening method(32) to rapidly and efficiently test the binding of radioligand to various hDAT mutants in detergent micelles. Briefly, plasmids encoding hDAT mutants were transfected into HEK293S cells immobilized on poly-D-lysine coated 96-well plates, and were incubated for 48 h. Post incubation, the cells were resuspended in a buffer containing MRS7292 and a radioactive cocaine analog, [^3^H]WIN35428. The plates were incubated for 30 min at room temperature before adding a mixture of Cu-YSi SPA beads and 4X detergent stock solution. Specific binding of [^3^H]WIN35428 to various hDAT mutants was measured using a SPA counter. If the mutation destabilized the transporter, the binding of [^3^H]WIN35428 and thereby the SPA signal, would be weak or nonexistent. A similar loss in the binding signal would ensue upon heat treatment(Fig 4).

**Fig 4:**
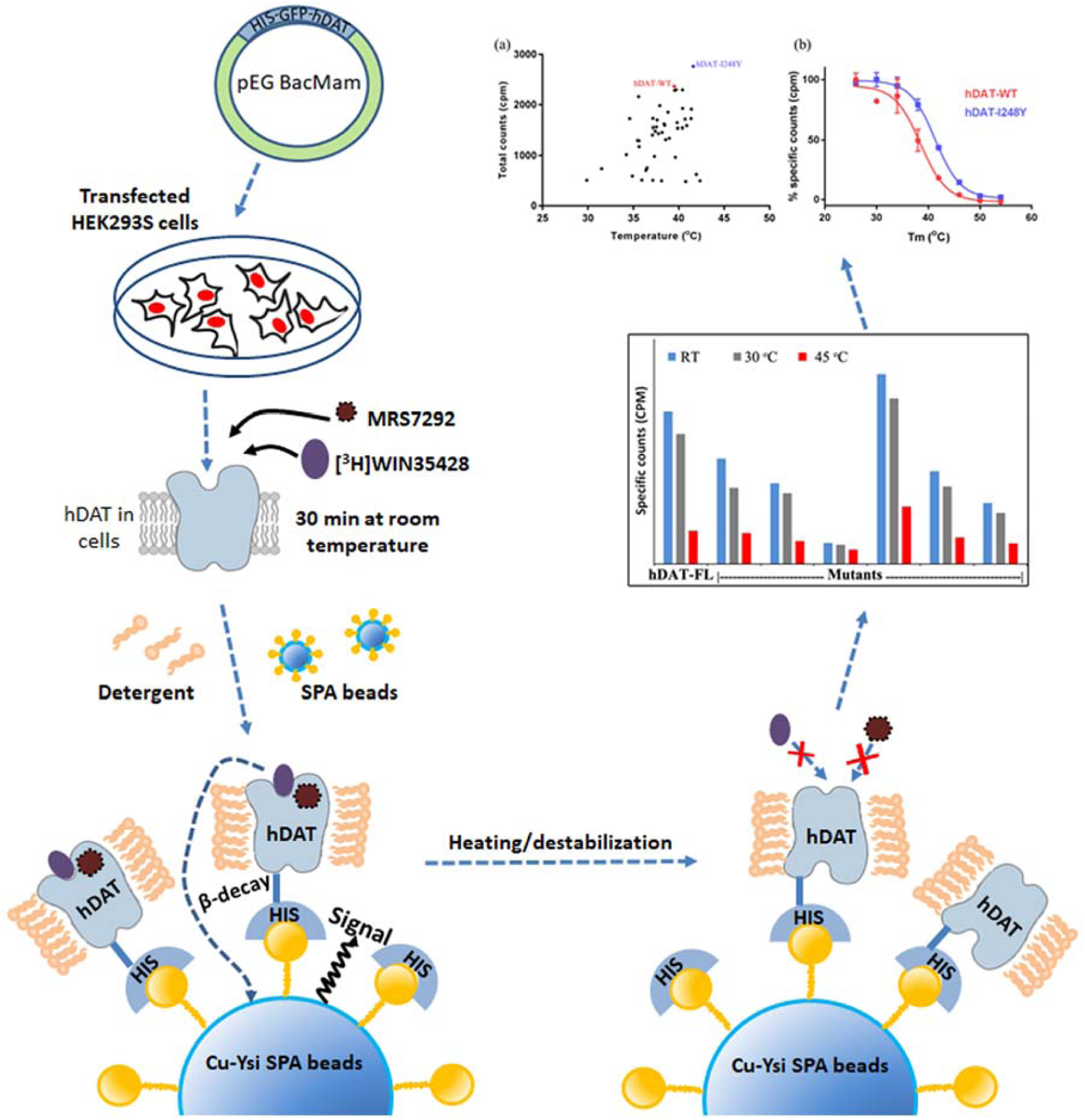
Experimental design for thermostability screening. Transiently transfected HEK293S cells expressing various mutants were resuspended in assay buffer consisting of MRS7292 and [^3^H]WIN35428. The suspension was incubated at room temperature for 30 mins to allow formation of ternary complex of hDAT and both the ligands. A mixture of detergent and SPA beads was added to end the incubation and begin protein solubilization. The counts from binding reaction were continuously monitored till they plateaued and the samples were subsequently subjected to heat treatment to measure the melting temperature. The decrease in specific counts with temperature was fit to Boltzmann sigmoidal equation to calculate *T*_*m*_ of mutants.

The temperature dependence of specific binding was estimated by subjecting the plates to incremental heat treatment. The mutants that displayed greater specific binding after heat treatment were retained for further analysis. The thermostable mutants were then expressed in HEK293S cells in 10 cm plates, and membranes containing hDAT variants were isolated. The isolated membranes were solubilized using detergent and radioligand binding was reexamined. A plot of apparent *T*_*m*_ versus total counts at room temperature was generated to more precisely evaluate the extent of thermostabilization. The I248Y substitution displayed both higher *T*_*m*_ and better expression than hDAT-FL(Fig 5a & 5b). We also identified a set of mutants that had marginally greater thermostability than hDAT-FL but displayed reduced expression. Finding only one promising thermostable mutant from a library of ∼200 mutants, despite the presence of MRS7292, indicates the extent of difficulty in thermostabilizing hDAT by conventional site-directed mutagenesis approach. Nonetheless, the successful mutant screening results encouraged us to attempt large scale purification of the I248Y variant, employing a similar detergent solubilization strategy as the one used in the SPA experiments.

**Fig 5:**
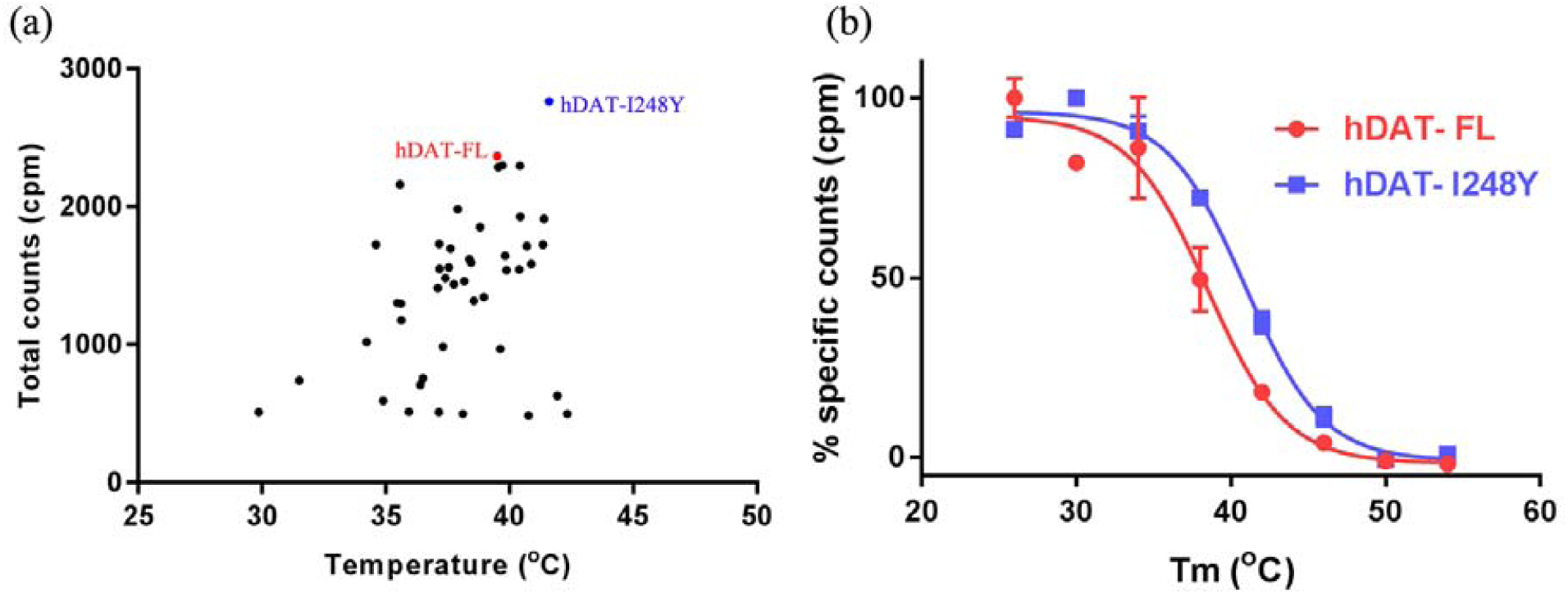
SPA based screening for thermostable mutants. (a) A scatter plot representing a subset of mutations(45 out of the total 200). The plot was generated by plotting total counts observed at room temperature versus apparent *T*_*m*_ of mutants, and(b) Boltzmann sigmoidal fit for melting temperature curves of hDAT-FL and most thermostable mutant. hDAT-FL is indicated in red, and hDAT-I248Y in blue. A representative of experiments(n=2 for panel a; n=5 for panel b) are shown, and the error bars in panel b denote standard errors of mean calculated from triplicate measurements.

### Purification of thermostable hDAT-I248Y variant

The SPA-based radioligand binding experiments indicated that occupancy of the allosteric site(MRS7292) and the central binding site(WIN35428) was important for stabilization of hDAT in detergent micelles. WIN35428 and GBR12909 are two high affinity ligands reported to bind central site on hDAT. We noticed that detergent solubilized hDAT-I248Y displayed a similar affinity to [^3^H]GBR12909(∼27 nM) and [^3^H]WIN35428(∼28 nM)(Fig 6a). Hence, we decided to use MRS7292 to saturate the allosteric site, and to saturate the central binding site we used non-radioactive GBR12909. Preliminary purification attempts comprised of solubilization of HEK293S cells expressing hDAT-I248Y in detergent containing buffer followed by affinity purification using TALON metal affinity chromatography and size-exclusion chromatography. However, the purified material was heterogeneous and was prone to non-specific proteolysis(Fig S3a). To circumvent this problem, we employed three strategies. First, we isolated cell membranes and resuspended them in a buffer containing MRS7292 and GBR12909 prior to solubilization(Fig S3b). Second, we replaced the GFP-tag in the expression construct with a Strep-tag II such that the resultant protein contained His_8_-StrepII-tag on the *N*-terminus enabling Strep-Tactin affinity purification and His-tag mediated SPA experiments(Fig S3c). Third, FSEC showed that the NΔ56-I248Y construct was less prone to aggregation and adventitious proteolysis(Fig S3d and S4; Fig 7), and thus we used this construct for large scale expression.

**Fig 6:**
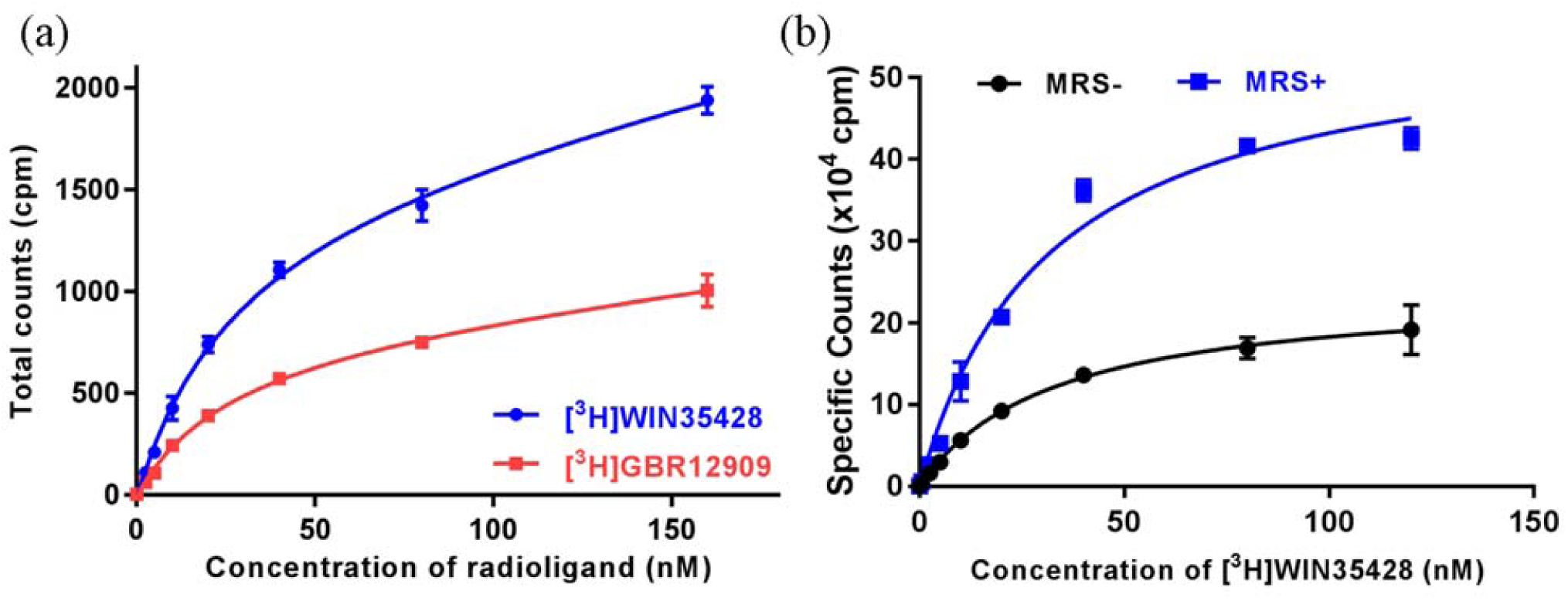
Radioligand saturation binding measurements. (a) Binding of [^3^H]WIN35428 and [^3^H]GBR12909 to purified hDAT-I248Y in presence of MRS7292, and(b) binding of [^3^H]WIN35428 to HEK293S cell membranes expressing hDAT-WT in presence and absence of MRS7292. A representative of experiments(n=1 for panel a; n=2 for panel b) are shown, and the error bars denote standard errors of mean calculated from triplicate measurements.

**Fig 7:**
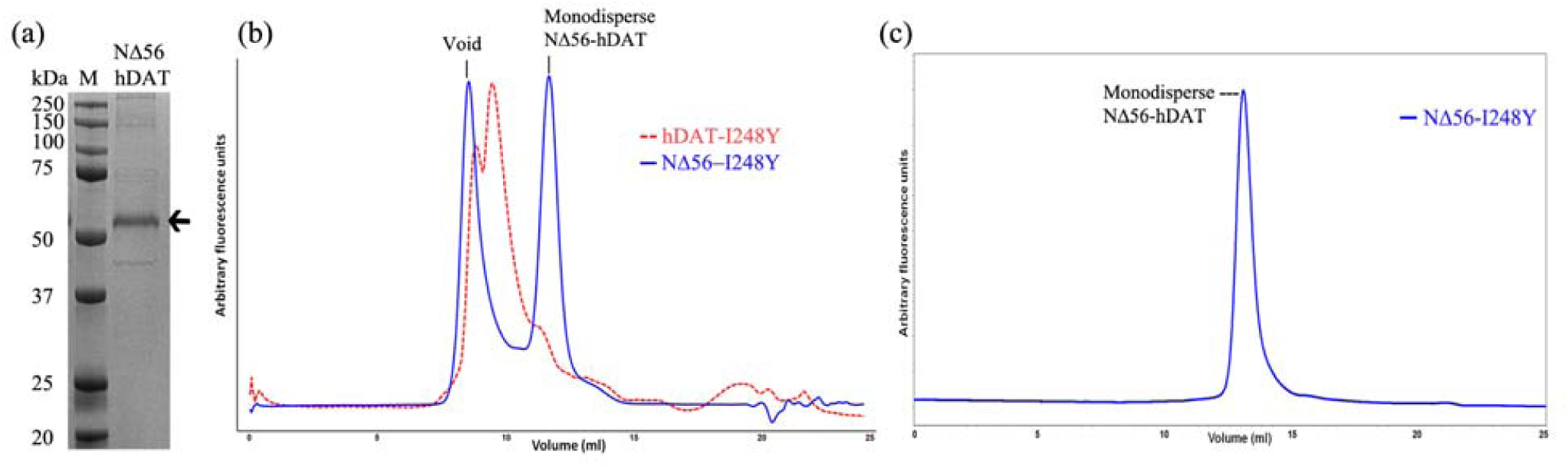
Purification of hDAT-I248Y. (a) 12% SDS-PAGE showing purified NΔ56-hDAT(indicated by arrow; M = protein molecular weight marker). M denotes protein molecular weight marker.(b) A comparison of preparative size-exclusion chromatograms of purified full length hDAT-I248Y(red) and its NΔ56 construct(NΔ56-hDAT). The protein eluted from Strep-Tactin affinity column(shown in panel a) was concentrated to 1mg/ml before loading onto Superdex S200 column for further purification, and(c) 0.5 μg of protein from the peak fraction of SEC step was rerun on Superose 6 column to analyze homogeneity. The intrinsic tryptophan fluorescence of hDAT was monitored in SEC and FSEC.

## Discussion

Neurotransmitter transporters are the major targets of psychostimulants. Molecular characterization of interactions between the psychostimulants and their targets can enable development of drugs with a lower potential for abuse. Despite sharing high sequence identity and a similar LeuT-structural fold, the pharmacology of neurotransmitter transporters is diverse, primarily because of the amino acid variations in the central binding site. The *Drosophila* dopamine transporter(dDAT) structure provided the initial structural basis for this observation. The residues in the central binding site of dDAT were more similar to hNET as opposed to hDAT or hSERT, an observation that explained the ability of dDAT to efficiently bind to ligands that are specific to hNET(nisoxetine and reboxetine), but not to hDAT(WIN35428)(22, 33, 38). In addition to pharmacological diversity, the sequence variation also determines the differences in the thermostability of these transporters. The mutations that conferred thermostability to dDAT and hSERT failed to exert a similar effect on hDAT. A major caveat of the hDAT mutant screening strategy when compared to dDAT or hSERT mutant screening procedure is the utilization of both a psychostimulant analog and an allosteric ligand(32, 33). The preliminary biochemical characterization of psychostimulant analog bound hDAT suggested that it is less stable when compared with anti-depressant bound dDAT or hSERT, underscoring the necessity for a modified mutant screening strategy.

The crystal structure of hSERT successfully identified an allosteric site in the extracellular vestibule of the transporter(39). It was hypothesized that the variations in the amino acid sequence of the allosteric site, especially in the EL6 region determined the specificity of allosteric site, and thus the allosteric ligands that modulated hSERT would not modulate hDAT or hNET. We focused our efforts to identify hDAT specific allosteric ligands with an assumption that a ternary complex of hDAT, psychostimulant, and an allosteric ligand, would be more stable than a binary hDAT-psychostimulant complex. We found reports which mentioned that certain rigid(N)-methanocarba-adenosine derivatives of A_3_ adenosine receptor agonists, containing a 5’-amide or 5’-ester group, a small *N*^6^-alkyl group and an C2-arylethynyl group(Fig S1a), specifically interact with hDAT(23, 24). These MRS compounds, bind to hDAT and enhance the binding of tropane derivatives to the central binding site. Variations in the functional groups attached to the methanocarba and adenine rings of the adenosine derivative could further modulate both the ligand binding and uptake activities. For instance, the derivatives containing a CH substitution at position 1 of the adenine ring showed no binding to hDAT(23, 24). We noticed that hDAT could retain its ligand binding activity in detergent micelles in presence of MRS compounds. Because variations in functional groups of MRS had an effect on binding, we presumed that they would also have an effect on stabilization of hDAT in detergent. Derivatives with CH substitution(instead of N) at position 3 of the adenine ring(MRS7173 and MRS7135) failed to preserve the ligand binding ability of hDAT upon detergent extraction. We determined the thermostability of hDAT in the presence of various MRS compounds that could preserve ligand binding activity using an SPA-based radioligand binding assay, and observed that derivatives containing an ester group attached to the methanocarba ring conferred moderately higher thermostability than those that contained an amide group. hDAT displayed highest *T*_*m*_ in presence of MRS7292(Fig S1). 5’-Methyl ester(MRS7292 and MRS7303) and 5’-ethyl ester groups(MRS7232 and MRS7301) enhanced the thermostability more effectively than a 5’-amide derivative(MRS5980). Based on the previous reports and the SPA results presented in this report, we believe that the derivatives containing 3-*deaza* modification of the adenine ring do not interact with hDAT efficiently.

We hypothesize that MRS7292 binds to hDAT at a site which is similar in location as that of the allosteric site on hSERT. In the hSERT-*(S)*-citalopram complex structure, two molecules of(*S)*-citalopram were detected, one at the central binding site and another at the allosteric site located in an outer vestibule(39). The dissociation of [^3^H](*R/S)-*citalopram from the central binding site of hSERT could be slowed in the presence of saturating concentrations of non-radioactive(*S)*-citalopram, which occupies the allosteric site and blocks ligand unbinding from the central binding site. In a similar fashion, diluting the hDAT-[^3^H]WIN35428 complex with binding assay buffer resulted in loss of counts with time, likely because of dissociation of the radioligand from the central binding site. Incorporation of 5 μM MRS7292 in the dilution buffer prevented the loss of binding(Fig 2b). However, unlike(*S)*-citalopram binding to hSERT, the binding of MRS7292 to hDAT is highly specific; it binds exclusively to the allosteric site of hDAT and does not bind at all to the central binding site, as evident by the lack of competition with [^3^H]WIN35428 to central binding site. Also, MRS7292 enhanced the B_max_ of binding by over 2-fold and marginally enhanced the affinity(Fig 6b). It is likely that MRS7292 binds to the allosteric site and sterically obstructs the exit of [^3^H]WIN35428 from the central site, an observation that encouraged us to design a screening strategy to identify mutants that could further stabilize the ternary complex of hDAT, MRS7292, and [^3^H]WIN35428.

We modified the screening strategy that was previously developed in the lab for screening hSERT mutants. We resuspended the transfected cells in a buffer containing MRS7292 and then incubated the mixture with [^3^H]WIN35428. We allowed the formation of the ternary complex prior to the addition of detergent, such that the bound ligands can shield hDAT from the destabilizing effects of detergent. Our mutant screening strategy resulted in the identification of one thermostable mutant, I248Y, which enhanced the *T*_*m*_ of hDAT by ∼ 2.4°C. We constructed a homology model of hDAT using the hSERT crystal structure(PDBID-5I6X) as template to identify the reason behind the apparent thermostability that is imparted by I248Y substitution. I248Y of TM4 lies in proximity of the residue F457 of TM9(Fig S2a). We hypothesize that substitution of I248 with aromatic residue such as Tyr promotes ring stacking interactions leading to greater stability. To test this hypothesis we generated a series of substitutions by replacing I248 with Trp, Phe, Leu, and Val and estimated the effect of these substitutions on thermostability. Only I248F showed enhanced *T*_*m*_, on par with I248Y. The I248W substitution, despite being aromatic residue incorporation like I248F or I248Y, failed to enhance the *T*_*m*_, likely due to the steric clashes introduced by the bulky indole ring of Trp residue. I1248L substitution appears innocuous when compared to I248V substitution(Fig S2b). Based on these observations, it appears that I248Y/F substitutions are better than the rest because they promote stacking interactions and are also sterically favorable. We believe that formation of ternary complex comprising hDAT, MRS7292, and [^3^H]WIN35428 stabilized hDAT in an outward facing conformation and enabled the identification of a thermostable variant of hDAT. Based on these observations we designed a purification strategy to isolate this ternary complex.

The detergent solubilization strategy employed in the purification protocol was designed to mimic the detergent solubilization in the SPA experiment, with one modification being the use of GBR12909 instead of WIN35428. Purification of hDAT required large quantities of ligands. GBR12909 had similar affinity to hDAT as that of WIN35428(Fig 6a), was readily available from commercial sources, and was much more affordable than WIN35428. Because, MRS7292 enhances binding of [^3^H]WIN35428 to hDAT, having non-radioactive GBR12909 bound at the central site allowed us to monitor the quality of purified material by performing a SPA assay in presence of saturating concentrations of MRS7292 and efficiently replacing GBR12909 with [^3^H]WIN35428.

The full length hDAT expression construct is susceptible to non-specific proteolysis, especially on the termini, because the extended intracellular *N*-(68 residues) and *C*-termini(40 residues) are populated with basic(Lys/Arg) residues that can act as sites for non-specific proteolysis(Fig 3 and S3). This is evident by a large free GFP peak in the FSEC based expression analysis(Fig S4). Both *N*-and *C*-termini are flexible and are believed to be involved in a plethora of non-covalent interactions with the innate lipids of the bilayer, the intracellular loops, and with a host of cytoplasmic proteins that are reported to play a crucial role in expression and trafficking of hDAT(40-43). Extraction of hDAT into detergent micelles will likely disturb such interactions, promoting further flexibility in the termini, leading to proteolysis and aggregation. To address these problems, we generated a series of constructs with deletions on the *N*-and *C*-termini of hDAT. We noticed that deletion of residues from C-termini, as opposed to *N*-termini, resulted in a decreased expression. The C-terminus is implicated in endocytic trafficking and internalization of hDAT(44). In addition, the crystal structure of dDAT elucidated that the C-terminus, through a series of hydrogen bond and cation-p interactions with IL1, forms a ‘C-terminal latch’ which could play a crucial role in the modulation of transporter activity(33).

To improve the quality of purification, in addition to modifying the expression construct boundaries, we also made changes to affinity purification procedure which resulted in enhanced purity. Removal of cytosolic proteins by first extracting membranes resulted in both a decrease in contamination as well as non-specific proteolysis(Fig S3). Mammalian proteins, by virtue of containing higher percentage of His residues, often result in impure purification when affinity resins such as Talon that bind to polyhistidine tag are used(45). Hence, further improvement in the purity of protein occurred upon changing the resin used in affinity chromatography from Talon to Strep-Tactin.

Despite the technological advances in the field of membrane protein expression and purification, the structural characterization of human neurotransmitter transporters is challenging because of the inability of these proteins to remain active or stable upon being extracted from the lipid bilayer. Identification of conditions that allow for extraction of an active form of transporter from the lipid bilayer is an essential prerequisite for biochemical and structural characterization. Here, we presented the development of a methodology for identification of transporter variants and conditions that aid in efficient purification of an active hDAT from the plasma membrane of HEK293S cells. We successfully purified a ternary complex containing hDAT, allosteric ligand, and the central binding site ligand. This complex, as well as the methods and hDAT constructs described here, will prove useful in the study of hDAT by structural and biochemical methods, ultimately aiding in the discovery of new small molecules that target the allosteric site(s), as well as the central binding site.

## Acknowledgements

We thank J. Coleman, F. Jalali-Yazdi, and other members of Gouaux lab for useful suggestions.

## Author contributions

V.N. and E.G. designed the experiments. D.K.T and K.A.J provided the MRS compounds. V.N. performed the experiments. V.N. and E.G. analyzed the data and wrote the manuscript.

## Supporting information

**Fig S1:**
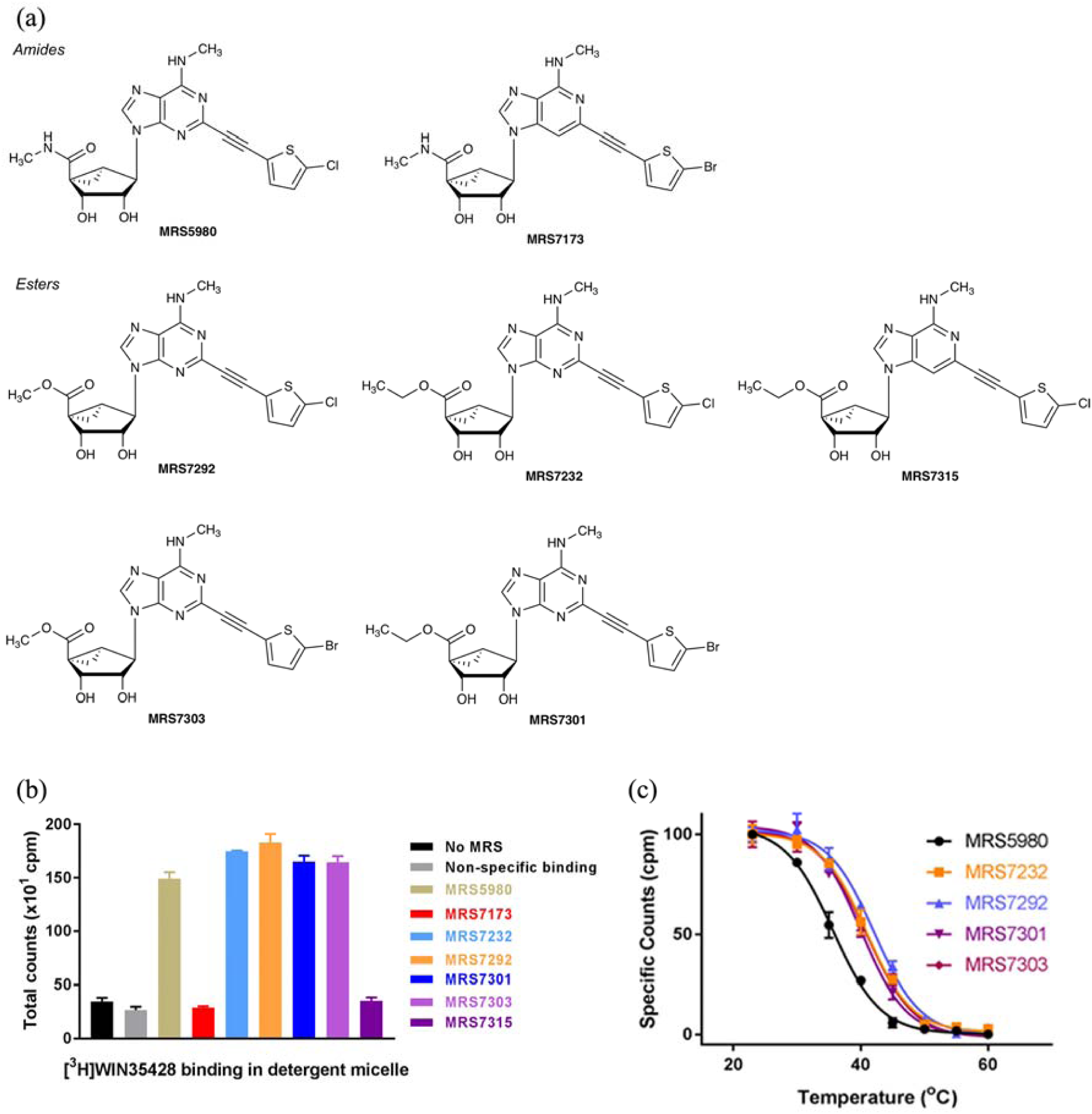
Effect of MRS compounds on stability of hDAT. (a) Chemical structures of MRS compounds tested in this study for their effect on hDAT thermostability,(b) A comparison of [^3^H]WIN35428 binding with and without MRS compounds. Compounds with *deaza* adenine moiety are ineffective in providing protection for hDAT against deleterious effects of detergent. (c) *T*_*m*_ of hDAT in complex with various MRS compounds. MRS7292 provides greatest enhancement in *T*_*m*_. The error bars represent standard errors of mean calculated from triplicate measurements of a representative experiment(n=2).

**Fig S2:**
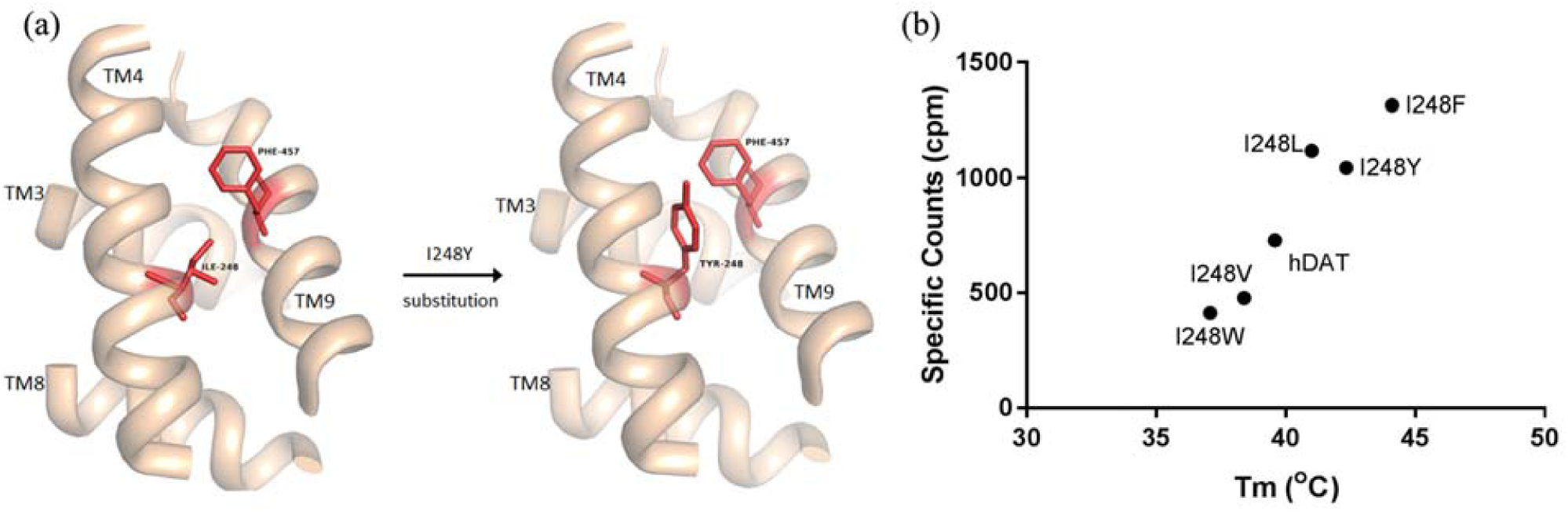
Rationale behind thermostability conferred by I248Y. (a) Substitution of I248 in TM4 with aromatic residues such as Tyr or Phe promotes stacking interactions with F457 in TM9 resulting in enhanced *T*_*m*_.(b) A substitution of a bulky or a tiny residue at I248 position disturbs this interaction and thus destabilizes the protein. These hypotheses are made on an hDAT model constructed using hSERT crystal structure as template.

**Fig S3:**
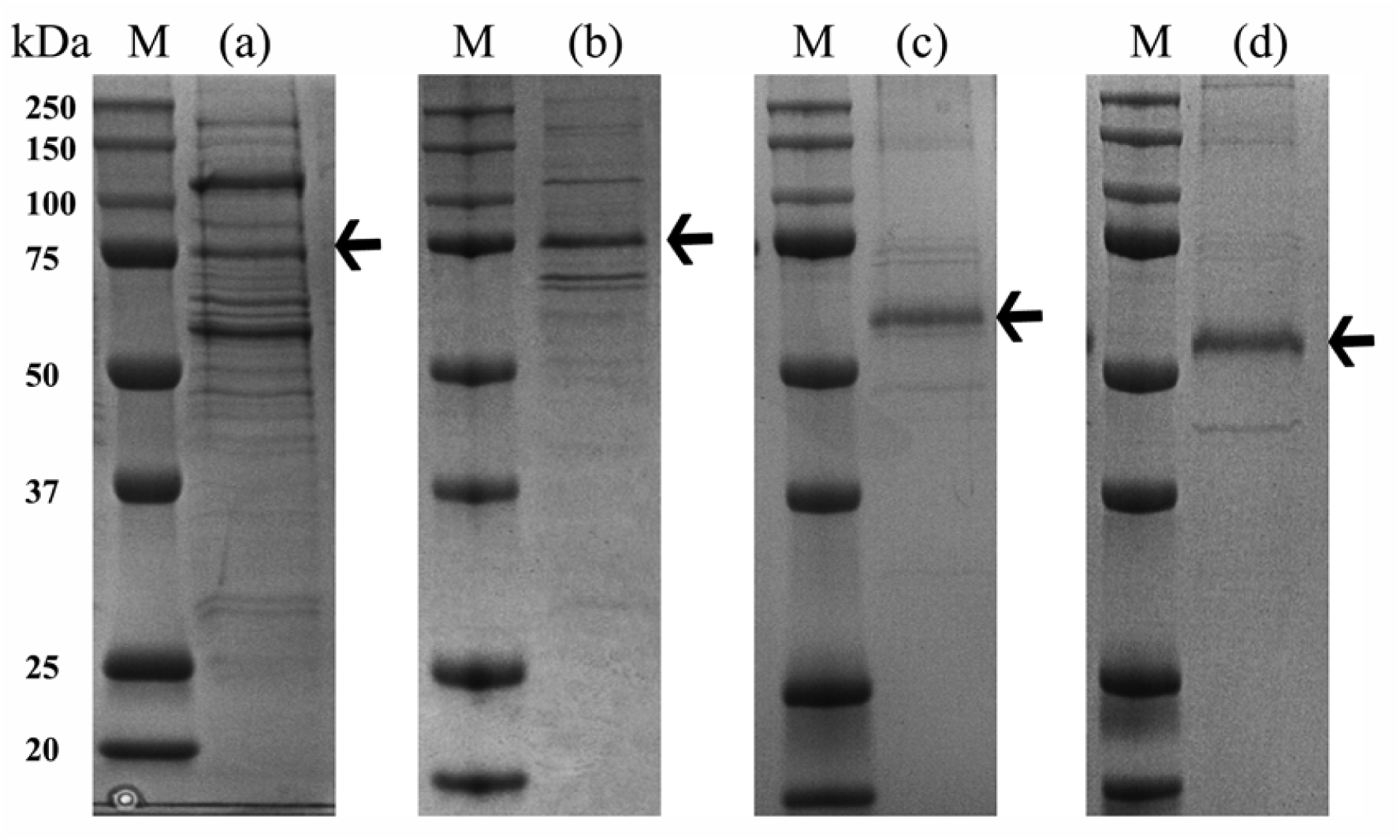
Comparative 12% SDS-PAGE analysis of optimization of purification of hDAT-I248Y. Purification(a) from whole cell solubilization using Talon affinity chromatography,(b) from solubilized membranes using Talon affinity chromatography,(c) using Strep-Tactin affinity chromatography, and(d) using NΔ56 expression construct and Strep-Tactin affinity chromatography. Position of hDAT in the lanes(a) &(b) were identified by in-gel GFP fluorescence. M denotes protein molecular weight marker

**Fig S4:**
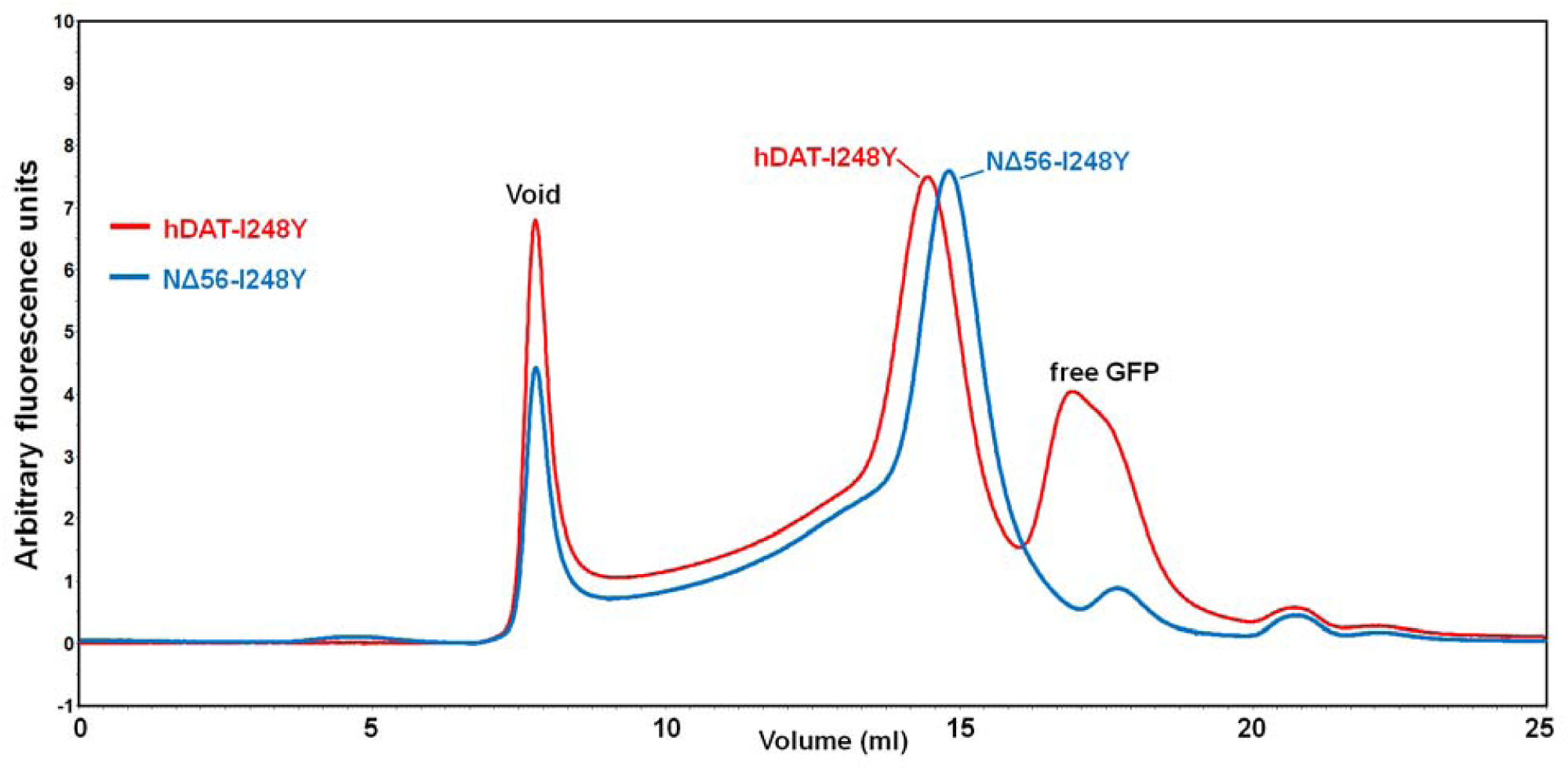
Comparison of expression profiles of full length and NΔ56 constructs of hDAT-I248Y. The *N*-terminal GFP fusion of hDAT-I248Y(red) and NΔ56-hDAT(blue) expression constructs are monitored by FSEC, using detergent solubilized cell lysate from HEK293S cells harvested 36 h post transfection. NΔ56 expression construct shows smaller void and free GFP peaks than full length. The free GFP is arising from non-specific proteolysis of *N*-termini.

